# Multivariate pattern analysis of brain structure predicts functional outcome after auditory-based cognitive training interventions

**DOI:** 10.1101/2020.09.06.283481

**Authors:** Lana Kambeitz-Ilankovic, Sophia Vinogradov, Julian Wenzel, Melissa Fisher, Shalaila S. Haas, Linda Betz, Nora Penzel, Srikantan Nagarajan, Nikolaos Koutsouleris, Karuna Subramaniam

**Author notes:** **corresponding author:** Lana Kambeitz-Ilankovic, PhD, Department of Psychiatry and Psychotherapy, University of Cologne, Germany, Kerpenerstr.62, D- 50937 Cologne, Germany.

## Abstract

**Background:** Cognitive gains following cognitive training interventions (CT) are associated with improved functioning in people with schizophrenia (SCZ). However, considerable inter-individual variability is observed. Here, we evaluate the sensitivity of brain structural features to predict functional response to auditory-based cognitive training (ABCT) at a single subject level.

**Methods:** We employed whole-brain multivariate pattern analysis (MVPA) with support vector machine (SVM) modeling to identify grey matter (GM) patterns that predicted ‘higher’ vs. ‘lower’ functioning after 40 hours of ABCT at the single subject level in SCZ patients. The generalization capacity of the SVM model was evaluated by applying the original model through an Out-Of-Sample Cross Validation analysis (OOCV) to unseen SCZ patients from an independent sample that underwent 50 hours of ABCT.

**Results:** The whole-brain GM volume-based pattern classification predicted ‘higher’ vs. ‘lower’ functioning at follow-up with a balanced accuracy (BAC) of 69.4% (sensitivity 72.2%, specificity 66.7%) as determined by nested cross-validation. The neuroanatomical model was generalizable to an independent cohort with a BAC of 62.1% (sensitivity 90.9%, specificity 33.3%).

**Conclusions:** In particular, greater baseline GM volume in regions within superior temporal gyrus, thalamus, anterior cingulate and cerebellum -- predicted improved functioning at the single-subject level following ABCT in SCZ participants.

## Introduction

Occupational and social functioning are impaired in patients with schizophrenia (SCZ) and are associated with a range of neural system and clinical impairments(1– 3). Cognitive training interventions (CT) can drive neural system changes (4,5) that are in turn associated with functional improvement (6–8). Previous studies have shown training-induced restoration of neural activation patterns in the medial prefrontal cortex and anterior cingulate cortex (mPFC/ACC) which were associated with improved performance on a reality monitoring task (4,9,10), and which in turn predicted durable gains in real-world social functioning 6 months later. We have also shown enhanced activation in the dorsal lateral prefrontal cortex, which was associated with improved performance on a working memory task, and predicted better occupational functioning at 6 months follow-up (11).

Although these findings are promising at the group level, it is clear that there is a large amount of inter-individual variability in neural system and functional response to various forms of CT. Previous group-based studies have shown that baseline structural anatomical integrity (12–14) is associated with greater responsiveness to CT, suggesting that certain individual neurobiological characteristics might determine who will benefit the most from this intervention, but individual-level predictions have not yet been demonstrated. In particular, prior research indicates that patients with schizophrenia (SCZ) show most prominent deficits in auditory processing, that contributed to higher-level cognitive impairments and poor functioning(15–17) Promisingly, we have also found that the most prominent gains in auditory/verbal functions were induced after auditory-based CT (ABCT) interventions (18–22) This work prompted us to investigate multivariate pattern analyses (MVPA) to identify baseline patterns in grey matter (GM) volume in patients with schizophrenia (SCZ) that predicted improved functioning after an auditory-based CT (ABCT) intervention, operating at the single subject level. Multivariate analyses of neuroanatomical brain properties have revealed high specificity of predicting improved functioning in Clinical High Risk (CHR) individuals with psychosis in single-site studies at the individual level (23,24), and have also shown remarkable multi-site generalizability (25). However, no study has yet examined the critical question of which structural features at baseline most predict responsiveness to ABCT, in terms of improving real-world functioning at the single subject level.

Group-based studies have shown that increased GM volume has been predictive of improved functioning in SCZ patients and also associated with stronger resilience to functional deterioration (1,12). Informed by these prior group-based studies and meta-analyses (1,12,13), we hypothesized that patients who had increased GM volumes at baseline, particularly in prefrontal, thalamic and temporal regions (1,12– 14) would show improved functioning at the single subject level in response to 40 hours of ABCT. We further explored the relationship between the decision values of the functioning classifier generated in SCZ patients and their clinical characteristics at baseline and at follow-up to test whether clinical symptom severity or medication dose were associated with individual level of functioning after 40 hours of CT. Finally, we investigated whether our original GM machine learning model was sufficiently generalizable to an independent training cohort that also underwent ABCT.

## Results

### Participant Characteristics

Table 1 summarizes the sociodemographic, clinical and cognitive characteristics of the two study samples. No significant differences with respect to age, premorbid IQ, years of education, illness duration and antipsychotic medication dosage (chlorpromazine equivalents) and GM volumes at baseline were found between lower and higher functioning SCZ in the original sample or in the IVS (p>0.05).

**Table 1.** Demographic and clinical data (baseline and post-training CT) for SCZ participants in Original and Independent Validation Samples, separated by their GAF score median split at the post-training timepoint. YoE = years of education, CPZ = chlorpromazine equivalent; GAF = Global Assessment of Functioning; QLS = Quality of Life Scale, PANSS = Positive and Negative Syndrome Scale

|  | Original Sample<br>(training time=40 h) |  |  |  | Independent Validation Sample (IVS)<br>(training time=50 h) |  |  |  | ORIG vs. IVS<br>for GAF<45 |  | IVS vs. ORIG<br>for GAF≥45 |  |
| --- | --- | --- | --- | --- | --- | --- | --- | --- | --- | --- | --- | --- |
| | GAF<45<br>(N=18) | GAF≥45<br>(N=18) | t/ $\chi^2$ | p | GAF<45<br>(N=10) | GAF≥45<br>(N=10) | t/ $\chi^2$ | p | t/ $\chi^2$ | p | t/ $\chi^2$ | p |
| age | 47.06(9.10) | 47.(8.99) | 0.05 | 0.95 | 38.45(13.19) | 41.0(11.84) | -0.45 | 0.65 | -2.08 | 0.05 | -1.52 | 0.14 |
| sex<sup>a</sup> | female=14 | female=14 | 0.00 | 1.00 | female=2 | female=3 | 0.60 | 0.44 | 9.80 | 0.02 | 5.08 | 0.02 |
| YoE | 13.39(1.69) | 13.22(1.86) | 0.28 | 0.78 | 12.82 (2.04) | 13.67 (3.12) | -0.73 | 0.47 | -0.81 | 0.42 | 0.46 | 0.64 |
| IQ | 103.56(11.28) | 100.82(12.30) | 0.68 | 0.49 |  |  |  |  |  |  |  |  |
| CPZ<sup>b</sup> | 265.5(129.40) | 299.2(212.49) |  | 0.92 | 398.71(354.95) | 290.06(188.2) | 0.82 | 0.42 |  |  | 0.45 | 0.64 |
| Illness Duration | 26.2(15.4) | 26.8(10.1) | 0.15 | 0.87 | 20.4(14.4) | 19.3(11.1) | 0.18 | 0.86 | -0.9 | 0.34 | 0.98 | 0.33 |
| GM volume | 645 (66.)6 | 602.8(63.9) | 343.35 | 0.06 | 656(84.0) | 674(65.1) | -0.53 | 0.60 | 0.372 | 0.71 | 2.71 | 0.01 |
| Training Intensity | 3.35(1.23) | 3.81(1.82) | -0.89 | 0.37 | n.a | n.a. |  |  |  |  |  |  |
| **Clinical Baseline** |  |  |  |  |  |  |  |  |  |  |  |  |
| PANSS Positive | 16.56(5.26) | 15.28 (5.83) | 0.69 | 0.49 | 21.09(6.07) | 19.56(6.41) | 0.54 | 0.59 | 2.12 | 0.04 | 1.74 | 0.09 |
| PANSS Negative | 16.67 (5.10) | 15.94 (6.11) | 0.38 | 0.70 | 21.91 (6.37) | 16.11(6.11) | 2.06 | 0.05 | 2.44 | 0.02 | 0.06 | 0.95 |
| PANSS General | 35 (7.51) | 30.61(9.11) | 1.57 | 0.12 | 41.82(8.26) | 36.0(4.2) | 1.24 | 0.23 | 2.28 | 0.03 | 1.27 | 0.21 |
| PANSS Total | 68.22(13.61) | 61.83(17.22) |  | 0.22 | 84.82(16.4) | 71.67(22.87) | 1.49 | 0.15 | 2.94 | 0.00 | 1.27 | 0.21 |
| GAF | 42.94(6.24) | 48.78(8.78) | -2.29 | 0.02 | 43.4(10.66) | 49.57(12.25) | -1.10 | 0.28 | 0.14 | 0.88 | 0.18 | 0.85 |
| QLS | 2.89(1.07) | 3.10(1.00) | -0.61 | 0.54 | 2.74(1.03) | 3.57(1.01) | -1.79 | 0.09 | 0.33 | 0.71 | 1.14 | 0.26 |
| **Clinical Post Training** |  |  |  |  |  |  |  |  |  |  |  |  |
| PANSS Positive | 17.33(4.50) | 13.50(3.33) | 3.70 | 0.00 | 19.36(4.31) | 17.78(5.56) | 0.71 | 4.82 | 1.19 | 0.24 | 3.12 | 0.00 |
| PANSS Negative | 16(4.74) | 14.06(5.59) | 1.12 | 0.26 | 23.36(5.46) | 15.00(7.48) | 2.88 | 0.01 | 3.83 | 0.00 | 0.37 | 0.71 |
| PANSS General | 34.61 (7.88) | 28.11(7.50) | 2.53 | 0.02 | 43.36 (10.97) | 37.89(11.95) | 1.07 | 0.3 | 2.49 | 0.01 | 2.61 | 0.00 |
| PANSS Total | 67.94(13.40) | 54.91(13.99) | 2.92 | 0.00 | 86.09 (14.64) | 70.67(23.58) | 1.79 | 0.09 | 3.41 | 0.00 | 2.23 | 0.03 |
| GAF | 40.2.(3.08) | 51.9(6.30) | -7.06 | 0.00 | 37.36 (7.29) | 56.56(7.37) | -5.82 | 0.00 | -1.47 | 0.15 | 1.71 | 0.09 |
| QLS | 3.0(1.10) | 3.28(1.06) | -0.76 | 0.44 | 3.01(0.79) | 3.94(1.05) | -2.24 | 0.04 | 0.14 | 0.97 | 1.5 | 0.15 |

However, significant differences in PANSS positive symptoms (t=3.70, p=0.001) and general PANSS symptoms (t=2.53, p=0.016) were observed between SCZ patients with lower vs. higher functioning in the original sample. In the IVS, SCZ patients with lower functioning had significantly more severe negative symptoms, compared to patients with higher functioning (t=2.88, p=0.04). Additionally, comparing patients with lower and higher functioning across two samples yielded significant differences in general PANSS symptoms (Table 1). Finally, the IVS consisted of more male participants (5.08, p=0.02), and the patients labeled as higher functioning exhibited significantly greater GM volume than patients from the original sample (2.71, p=0.01).

### Performance of Original Classification Model

SCZ subjects with GAF<45 had significantly lower functioning than GAF≥45 both at baseline and after ABCT (t=6.67, p<.0001; t=7.13, p<.0001). Due to the heterogeneous response to ABCT we did not find a significant difference in GAF between baseline and post-training at the group level (t=.26, p=.80). However, we found that SCZ subjects with GAF≥45 after ABCT showed significant improvement in GAF scores from baseline (t=2.2, p=.05). Based on this ground GAF of ≥45 was determined as the most accurate median-split score for the sMRI classifier to determine the structural features at baseline that best predicted which individuals would show significantly improved and better functioning after ABCT, compared to baseline and compared to SCZ patients with GAF<45.

The sMRI GM classifier correctly discriminated SCZ patients in the original sample with higher functioning from lower functioning after the CT intervention with a cross-validated BAC of 69.4%, Sensitivity= 72.2%, Specificity=66.7%, NPV=70.6% and NND of 2.6. The permutation analysis showed that the classification models produced by the binary GAF classifier in response to ABCT were significant at p=.00.

Inspection of the mean feature weights generated within the CV framework revealed that the classification of the higher functioning from lower functioning patients in response to the CT intervention was driven by increased baseline GM volumes in primarily temporal regions (i.e., in bilateral superior and inferior temporal regions, including ventral visual word form area, and parahippocampal gyri), thalamic and frontal regions in the anterior cingulate cortex, as well as increases in the posterior cingulate cortex and cerebellum (Figure 1). Though the GM pattern was mainly characterized by baseline volume increases in the higher functioning patients, the lower functioning group also showed some volume increase in primarily motor (i.e., premotor cortex and supplementary motor area) and caudate regions within the basal ganglia (Figure 1).

**Figure 1.**
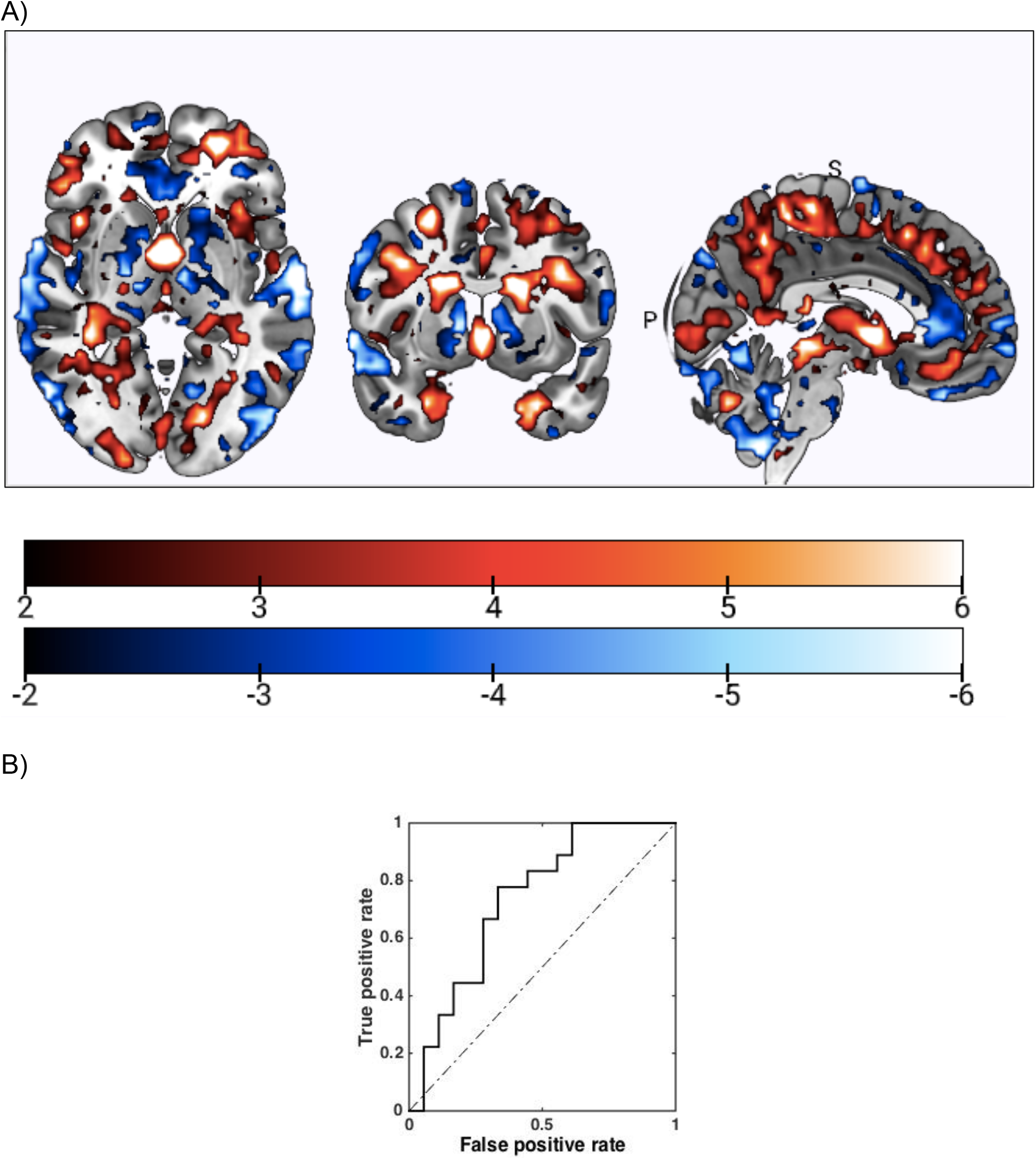
Structural MRI-based Classifiers Predict Functional Response to Auditory-based Cognitive Training. A) The reliability of predictive pattern elements in significant outcome classification models was measured in terms of a Cross-Validation Ratio (CVR) map (CVR = mean(w) / standard error(w), where w = the normalized weight vectors of the SVM models. Warm color scales indicate increased vs. decreased GM volume in the SCZ subsample with post-training GAF<45 vs. GAF≥45. Cool colors indicate increased vs. decreased GM volume in the GAF**≥**45 vs. GAF<45 subsamples. B) Receiver-Operator-Curve of the class probability values obtained from the trained L1-MVLR model in unseen SCZ persons, as determined by nested cross-validation.

Although the MRI classification model provided accurate estimates (e.g. BAC=69.4%) of correctly discriminating higher vs. lower functioning in response to the ABCT, we also wanted to ensure that the MRI-based classifiers did not predict generic baseline variations in global functioning that were not specific to the ABCT and may have confounded response to the ABCT. To investigate this possibility further, we replaced the GAF post CT functioning labels of the SCZ patients at follow-up with the respective classification labels derived from the baseline GAF scores and repeated the SVM analyses with the same machine learning pipeline as described previously. The MRI classifier differentiated lower from higher functioning SCZ with a non-significant classification at chance level with a BAC 44.4% (sensitivity 44.4%, specificity 44.4, NPV 44.4%). These results indicate that the individual structural features were therapeutically specific to predicting response to the CT intervention, rather than general baseline functioning levels.

### OOCV Model Performance

We next applied the original GM classification model to the IVS to predict follow-up functioning after the ABCT intervention in order to test whether the original GM classification model would generalize to the IVS. The model was able to successfully discriminate lower from higher functioning SCZ patients in the IVS at post ABCT sufficiently above chance with a BAC of 62.5, Sensitivity 90.9 % and Specificity 33.3%, NPV of 75.0 and NND of 4.1.

### Decision Scores and Correlational Analysis

The decision values of the discriminative GM signature for lower vs. higher functioning after ABCT were not associated with the baseline functioning levels (r=−3.02, p=0.07), indicating that the prediction of response to CT was not confounded by baseline patterns but was specific to predicting response to the intervention.

We investigated the relationship between decision scores in both the original and OOCV models with PANSS symptoms as well as decision scores with sex, in order to exclude the possibility that the original classification was biased by respective differences within and between the two samples. None of the associations yielded significant results (p>0.05).

We also correlated decision scores with antipsychotic medication dosage as assessed via CPZ equivalents, and found significant associations (r=0.42, p<0.02) in the original sample, suggesting that higher doses of medication were associated with better functioning after the CT intervention. No such correlation between decision scores and CPZ medication was observed in the IVS (r=0.08, p=0.53).

## Discussion

This is the first study to apply MRI-based machine learning to predict individual functional responses to an intensive course of ABCT in chronically-ill SCZ patients. The original classification model provided accurate estimates of 69.4% in correctly discriminating higher vs. lower functioning after the ABCT. Importantly, the MRI-based classifiers did not predict baseline variations in functioning, indicating that the individual structural features were therapeutically specific to predicting response to the CT. The original classification model generalized to an independent validation sample (IVS) with an accuracy of 62.5%. These results confirm that GM volumes have high predictive specificity for individual therapeutic functional response to our ABCT intervention in SCZ patients. Multivariate pattern analyses methods thus have the potential to use neuroanatomical biomarkers to predict functional response to therapeutic interventions at the individual level (26,27).

However, due to the heterogeneous response to ABCT, we did not find a significant difference in GAF between baseline and post-training at the group level, and thus used a median split of GAF score of 45 which significantly differentiated SCZ with high versus low functioning both at baseline and after the CT intervention. Although at a group level, we did not find a significant difference in GAF at post-training compared to baseline, this proved to be a strength rather than a limitation of the machine learning classifier as our findings revealed that it was still sensitive to subtle changes in GAF that allowed the classifier to accurately predict which individuals showed gains in individual functional responses after the ABCT compared to baseline.

Two prior studies have shown that, at the group level, SCZ patients with higher GM volumes at baseline showed a stronger response to cognitive training (1,12), GM volumes have also been shown to increase in response to cognitive training interventions (13). However, group-level analyses cannot take into account the substantial individual neuroanatomical heterogeneity that occurs at the individual level in SCZ (28). The goal of precision-medicine is to select and adapt therapeutic approaches based on each patient’s individual neural and clinical characteristics (29,30). In order to take into account individual neuroanatomical heterogeneity at baseline and reliably validate the origin of the predictive information, we replaced patients’ GAF scores at post ABCT with their baseline scores and repeated the sMRI GM analysis. Strikingly, we were not able to find significant GM patterns that successfully discriminated patients with lower from higher functioning at baseline. Moreover, the decision values of the discriminative GM signature for lower vs. higher functioning were not associated with baseline functioning levels. These results indicate that the prediction of functional response to CT was not confounded by baseline functioning.

Accurate discrimination of participants with higher functioning after the CT intervention (GAF scores of ≥45) was specifically shown by GM volume increases in superior temporal gyrus (STG), ventral visual form areas, thalamus and parahippocampal gyri. Longitudinal studies indicate progressive decreases in STG volume after the first psychotic episode, and this neuroanatomical abnormality is consistently reported in people with established SCZ (33,34). These results are consistent with our prior group-based studies showing the functional importance of the STG during ABCT, and its responsiveness to our ABCT interventions (18,22). We have specifically shown increased recruitment of the primary auditory cortex and the prefrontal cortex mediating auditory learning after ABCT (22). Additionally, the results from our recent study indicate that at baseline, even chronically-ill SCZ suffering from hallucinations were able to recruit the visual ventral word form area, which correlated with auditory and verbal working memory (9) Our data suggest that an intact GM reserve particularly in the STG at baseline drives plasticity to auditory cognitive training interventions, and is likely to predict which SCZ patients receive most benefit from auditory training interventions.

We have previously shown at a group level, that the intact structure of thalamic-prefrontal regions was also an important determinant of successful responsivity to cognitive training interventions in SCZ (14). Additionally, Ramsay et al showed that cognitive training was associated with increased thalamic-prefrontal activity and connectivity such that improved recruitment after cognitive training became correlated with overall improved cognitive functions (33). Importantly, previous studies have shown that the ACC/mPFC plays a critical role in supporting higher-order cognitive control functions that are important for conflict resolution and reality-monitoring functions (34–36). We have previously shown that increased recruitment of the ACC/mPFC induced by our cognitive training, correlated with successful reality monitoring performance that generalized to improved long-term social functioning (9). These prior studies support the data in the present study, in which we found increased GM volume in the ACC that predicted better functioning after ÁBCT.

Interestingly, we also found that SCZ who showed greater GM volume increases in the cerebellum at baseline, also revealed better overall functioning induced by ABCT. The cerebellum is important for mediating sensory prediction-errors for updating an internal model of implicit learning and action-outcome behaviors that are fundamental for improving real-world functioning in schizophrenia(37–39). These data are consistent with the neuroplasticity principles of our ABCT intervention, which specifically trains SCZ to improve auditory detection, temporal integration, prediction-error and learning, that have shown to directly contribute to higher-level functioning (18,40). In summary, SCZ patients who exhibit relatively increased GM structural volumes at baseline in STG, thalamus, ACC and cerebellum, in particular, may possess the needed neurological infrastructure to maximally benefit functionally from intensive ABCT. Together, our present findings are consistent with these prior meta-analyses and group-based studies, indicating that the individual predictive value of recruitment of regions particularly in the cerebellum, STG and thalamic-prefrontal areas are important and critical targets for cognitive training interventions.

It must also be noted that we also found that accurate discrimination of SCZ participants with lower functioning (i.e., GAF score of <45) was characterized by GM increases in premotor and basal ganglia regions. Aberrant connectivity in the motor system and disturbances in motor behavior have been observed in SZ patients with lower functioning (41,42). Basal ganglia volume increase (hypothesized to be due to striatal hyperdopaminergia) has also been shown in both medicated and antipsychotic-naive patients in meta-analytic studies (39) concurrent with motor disturbances as a one of the central clinical features of SCZ.

Some studies have raised the question as to whether GM loss or increase can be attributed to cumulative exposure to antipsychotic medications, rather than to aberrant neural developmental processes (43). Importantly, the decision values of our SVM analysis that accurately predicted higher functioning in response to the CT intervention, showed a significant relationship with medication dosage. Specifically, SCZ patients who had a higher medication dosage at baseline as well as lower positive symptoms, also had better functioning following the intervention. Taken together, these findings suggest that structural features together with medication dosage provide useful determinants of individual functional responsiveness to cognitive training interventions at the single subject level.

The main limitation of the present study is that the findings here do not account for the heterogeneity associated with additional neurophysiologic, environmental, and genetic factors that play a role in the response to ABCT. In order to develop a more robust and definitive predictive model, future studies will require: 1) a wide variety of behavioral and neurophysiologic data analyzed in a multivariate fashion to develop more accurate predictive biomarkers, so that meaningful signals are less likely to be lost due to noise from highly variable and heterogeneous metrics (44); 2) larger study samples from a wide variety of multisite studies, in order to provide more extensive geographical generalizability; 3) participants with a range of illness durations. The participants in our study had, on average, been ill for more than 20 years, limiting the generalizability of our findings to only older people with chronic illness who are also likely to manifest more severe symptoms.

In conclusion, with our whole-brain MVPA analyses, we have identified a structural MRI fingerprint associated with preserved GM volumes within particular regions in the STG, thalamus, ACC and cerebellum, that predicted improved functioning following an ABCT intervention, and that serves as a model for how to facilitate precision clinical therapies for SZ based on imaging data. Future studies should investigate if the individuals with greatest GM loss in these regions (who may also have the greatest vulnerability for more subsequent and severe psychotic episodes) can benefit from enhancements to CT that might include more intensive and integrative therapies of combined pharmacotherapy, cognitive training (45) and neuromodulation (46). If identified early in young adulthood, cerebellum-temporal-thalamic-prefrontal GM loss may reflect an important opportunity to provide early and intensive interventions to mitigate and reduce the impact of future and more severe psychotic episodes on functioning (47,48).

## Methods

### Participants and Procedure Overview

Two independent samples of SCZ participants who had structural imaging data were drawn from two larger clinical trials of that have been previously reported (ClinicalTrials.gov Identifier: NCT02105779)(49,50) and (ClinicalTrials.gov NCT00312962) (11). As is customary in predictive analytics, the MVPA model was constructed from the first set of subjects (the original sample, N=44) and then applied to a different set of subjects (the independent sample, N=23) using an out of cross-validation (OOCV) approach. This process produces an unbiased estimate of the method’s predictive accuracy on new individuals rather than merely fitting the current study population (51). The study was carried out in accordance with The Declaration of Helsinki, and reviewed by the Institutional Review Board at the University of California, San Francisco.

All SCZ subjects were recruited from community mental health centers and outpatient clinics. Inclusion criteria were: Axis I diagnosis of schizophrenia, schizoaffective disorder, or psychosis not otherwise specified (NOS) (determined by the Structured Clinical Interview for DSM-IV [SCID])(52). All participants provided written informed consent and then underwent structural imaging, clinical and cognitive assessments at baseline and after the CT intervention. Participants with poor signal-to-noise ratio in their neuroanatomical images were excluded from the final analyses for the original (N=5) and IVS (N=1) samples. Three participants from the original sample and two participants from the IVS cohort did not complete functioning assessments at the follow-up time point. An overview of our procedures can be found in Figure 2. Demographic characteristics of the two samples are presented in Table 1.

**Figure 2.**
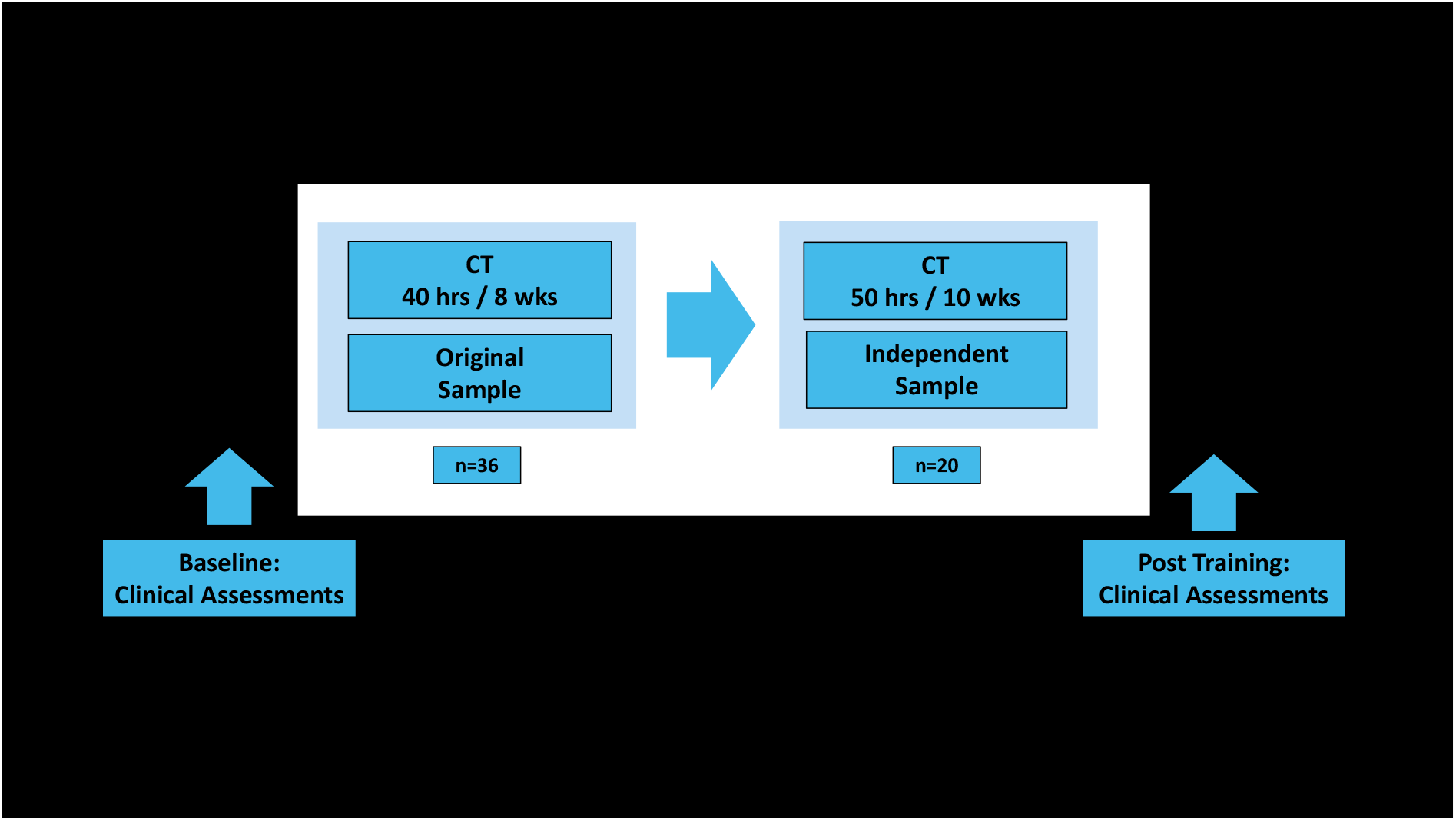
Training Design of the Original and Independent Validation Samples. The machine learning support vector model reliably predicted GAF≥45 vs. GAF<45 in SCZ participants in response to the auditory-based cognitive training (CT) at the single subject level in both samples.

### Auditory-based CT Intervention

The design of our neuroscience-informed computerized CT intervention is based on 3 decades of research into known mechanisms of neural plasticity, which have been shown to increase neuronal activity, synaptic connectivity and neuronal fiber integrity (53,54). In particular, research has documented the neural plasticity of cortical responses as an individual acquires new perceptual and cognitive abilities (55). A rich body of work shows that improved functioning after our cognitive training regimens specifically results from neuroplasticity (defined as neural structural and functional changes induced by the CT intervention). Complete details of the ABCT exercises can be found here (http://www.positscience.com/our-products/brain-fitness-program). Briefly, in the original sample performed auditory exercises for 1 hr a day for a total of 40 hours of training and the IVS sample performed 50 hours of training over the course of ∼10 weeks. In the exercises, patients were driven to make progressively more accurate discriminations and temporal integration about the spectro-temporal fine-structure of auditory stimuli under conditions of increasing working memory load under progressively briefer presentations, and to incorporate and generalize those improvements into working memory rehearsal and decision-making. The auditory exercises were continuously adaptive: they first established the precise parameters within each stimulus set required for an individual subject to maintain 80% correct performance, and once that threshold was determined, task difficulty increased systematically and parametrically as performance improved(20).

### Clinical and Functional Outcome Assessments

The Structured Clinical Interview for DSM-IV Axis I Disorders (56) was administered at baseline to all participants. In both the original GM machine learning and the IVS samples, Global Assessment of Functioning (GAF) Scale of the DSM-IV (57), and Quality of Life Scale (58) were used to assess functioning and the Positive and Negative Syndrome Scale (PANSS)(59) was used to assess severity of clinical symptoms at baseline and after the CT intervention. Functional assessments such as the Global Assessment of Functioning (GAF) quantify how much a person’s symptoms affect his or her real-world day-to-day life on a scale of 0 to 100.

### Machine Learning Strategy

Following our aims, we employed a nested cross-validated machine learning pipeline to evaluate the sensitivity of GM volumetric features at baseline to predict GAF functional response to cognitive training at a single subject level, using a median split strategy with the machine learning analyses delineated below (24,25).

### Determining Median Split

GAF scores were used to determine labels of lower vs. higher functioning after CT by setting a median split as a cut-off. GAF ≥45 determined selection criteria for patients with higher functioning (n=18) whereas GAF<45 determined selection criteria for patients with lower functioning (n=18). In our OOCV analysis, the median split cut-off was identical, with 10 patients with lower and 10 with higher functioning in the IVS. The rationale for determining a GAF score of 45 as a cut-off to split higher functioning SCZ patient from lower functioning SCZ patients was based on several factors: 1) At baseline, SCZ subjects with GAF<45 had significantly lower functioning from GAF≥45; 2) SCZ subjects with GAF≥45 after auditory CT showed significant improvement in GAF scores from baseline; and 3) In our chronic SCZ sample, a GAF score of 45 represents moderate impairment in functioning, and was also the most representative level of functioning that the chronic SCZ patient population experience (e.g. range of GAF scores in the chronic SCZ patients was from 29-67 in the original sample and 23-70 in the OOCV sample).

### Machine Learning Analysis Pipeline

The in-house machine learning platform NeuroMiner, version 1.0 (25) was used to set up a machine learning analysis pipeline in which the individual classification ability (‘higher’ SCZ functioning-’lower’ SCZ functioning); obtaining GM volume was performed for the prediction target GAF. To strictly separate the training process from the evaluation of the predictor’s generalization capacity and prevent the leakage of information and overfitting, the pipeline was completely embedded into a nested cross-validation framework (nCV) (31) with a 10-by-5 cross-validation (CV) structure for both inner (CV1) and outer (CV2) cycles. Only the inner CV1 training cycle was used for the implementation of the preprocessing steps (e.g. scaling, hyperparameter optimization) which does not occur again on the outer cycle (CV2), but CV2 serves exclusively the purpose to measure the models’ generalizability to new, unseen data. This procedure was applied to each fold for each permutation combination independently.

This analysis chain was applied to the outer CV cycle determining the patients’ classification of higher vs. lower functioning through majority voting determining balance accuracy (BAC) of the Support Vector Machine (SVM) model. Statistical significance of individual classifiers was assessed through permutation testing, with α=0.05 and 1000 permutations. Further information on this approach can found in in Koutsouleris et al.(60).

### Statistical analysis

Sociodemographic differences between groups were examined using analysis of variance (ANOVA) for parametric data, and by χ^2^ test for non-parametric data, as implemented in Jamovi for Windows (0.9.5.12). Furthermore, potential interactions between subjects SVM functioning decision scores and their (1) GAF levels at baseline, (2) antipsychotic medication doses (in chlorpromazine equivalents), and (3) clinical symptoms at baseline and follow-up were assessed by means of correlational analysis (Pearson’s r). Significance was defined at P < 0.05.

## Acknowledgments

This research is supported by the Brain and Behavior Research Foundation Young Investigator Award grants (NARSAD: 17680 and 28188), and NIMH K01 grant (K01MH105615) and R01 grants (R01MH122897) to Karuna Subramaniam, as well as an NIMH R01 grant (R01MH82818) to Sophia Vinogradov. We are also thankful to Brain and behavior Research Foundation for supporting work of Lana Kambeitz-Ilankovic through Young Investigator Award No° 28474).

## Author Contributions

LKI, KS and NK conceptualized the paper. KS, MF and SV oversaw data collection and project development. LKI and KS were responsible for statistical analyses. LKI, KS and SV drafted the manuscript. LKI, KS and SN provided data interpretation. SH, JW, LB, and NP assisted with neuroimaging procedure, statistical analyses and data interpretation. All authors revised and agreed upon the final version of the manuscript.

## Notes

### Competing Interest Statement

The authors have declared no competing interest.

