## Supplemental methods for "Multivariate pattern analysis of brain structure predicts functional outcome after auditory-based cognitive training interventions"

### Neuroimaging Protocol

In the original sample, imaging was performed on a 3 Tesla Siemens Prisma MRI scanner with 64- and 20-channel head and neck coils at the Neuroscience Imaging Center at University of California San Francisco.
In the IVS, high-resolution anatomical images were acquired from each individual on a 3 T General Electric Signa LX 15 scanner, utilizing 3D magnetization prepared rapid gradient echo MRI (160 1-mm slices; FOV = 256 mm, matrix = 256 × 256, TE = 2 ms, TR=7ms, flip=15).

### sMRI Processing Pipeline

The manual of the CAT12 toolbox, version r>1200 (http://www.neuro.uni-jena.de/cat12/CAT12-Manual.pdf) details the processing steps applied to the structural images (Koutsouleris et al. 2018). These steps consist of:

(1) A 1st denoising step based on Spatially Adaptive Non-Local Means (SANLM) filtering.
(2) An Adaptive Maximum A Posteriori (AMAP) segmentation technique, which models local variations of intensity distributions as slowly varying spatial functions and thus achieves a homogeneous segmentation across cortical and subcortical structures.
(3) A 2nd denoising step using Markov Random Field approach which incorporates spatial prior information of adjacent voxels into the segmentation estimation generated by AMAP(2).

(4) A Local Adaptive Segmentation (LAS) step, which adjusts the images for white matter (WM) inhomogeneities and varying gray matter (GM) intensities caused by differing iron content in e.g. cortical and subcortical structures. The LAS step is carried out before the final AMAP segmentation.
(5) A Partial Volume Segmentation algorithm that is capable of modeling tissues with intensities between GM and WM, as well as GM and cerebrospinal fluid (CSF) and is applied to the AMAP-generated tissue segments.
(6) A high-dimensional DARTEL registration of the image to a MNI-template generated from the MRI data of 555 healthy controls in the IXI database (http://www.braindevelopment.org). The registered GM images were multiplied with the Jacobian determinants obtained during registration to produce GM volume maps.
(7) GM images were smoother with a Gaussian smoothing kernel with a 4mm full width at half maximum (FWHM).

The Quality Assurance framework of CAT12 was used to empirically check the quality of the GMV maps. By computing the correlation of each image to all other images, taking the original and independent sample separately, we removed two images whose correlation exceeded 2 standard deviations from the sample mean due to MRI artifacts.

**Machine Learning Preprocessing Parameters**

We regressed out the effect of the total GM volume by entering the values as a covariate. In the inner CV loop, zero-variance features were pruned. Then, a dimensionality reduction procedure was applied through Principal Component Analysis (PCA). In order to minimize the generalization error, PCA models were trained with a limited number of Principal Components (PC) in the CV1 training data (80 PC) and subsequently scaled (from 0-1). The C hyperparameter was optimized in the range from 0.0156 to 16.
Model performance was measured using Sensitivity, Specificity, Balanced Accuracy (BAC), Positive Predictive Value (PPV), Negative Predictive Value (NPV), and Number Needed to Diagnose (NND), based on the class membership probability scores generated through majority voting in the outer CV cycle.
